# Efficient Merging of Genome Profile Alignments

**DOI:** 10.1101/309047

**Authors:** André Hennig, Kay Nieselt

## Abstract

**Motivation:** Whole-genome alignment methods show insufficient scalability towards the generation of large-scale whole-genome alignments (WGAs). Profile alignment-based approaches revolutionized the fields of multiple sequence alignment construction methods by significantly reducing computational complexity and runtime. However, WGAs need to consider genomic rearrangements between genomes, which makes the profile-based extension of several whole-genomes challenging. Currently, none of the available methods offer the possibility to align or extend WGA profiles.

**Results:** Here, we present `GPA`, an approach that aligns the profiles of WGAs and is capable of producing large-scale WGAs many times faster than conventional methods. Our concept relies on already available whole-genome aligners, which are used to compute several smaller sets of aligned genomes that are combined to a full WGA with a divide and conquer approach. To align or extend WGA profiles, we make use of the SuperGenome data structure, which features a bidirectional mapping between individual sequence and alignment coordinates. This data structure is used to efficiently transfer different coordinate systems into a common one based on the principles of profiles alignments. The approach allows the computation of a WGA where alignments are subsequently merged along a guide tree. The current implementation uses `progressiveMauve` (Darling *et al.*, 2010) and offers the possibility for parallel computation of independent genome alignments. Our results based on various bacterial data sets up to several hundred genomes show that we can reduce the runtime from months to hours with a quality that is negligibly worse than the WGA computed with the conventional `progressiveMauve` tool.

**Availability:** `GPA` is freely available at https://lambda.informatik.uni-tuebingen.de/gitlab/ahennig/GPA. `GPA` is implemented in Java, uses `progressiveMauve` and offers a parallel computation of WGAs.

## 1 Introduction

Whole-genome sequencing (WGS) has become increasingly affordable through the continuous developments of next-generation sequencing (NGS) technologies. WGS for example is now routinely conducted in a clinical context to monitor pandemic bacterial outbreaks based on the sequencing of different isolates. Single nucleotide polymorphisms (SNPs) between isolates and a reference genome can help to understand and reconstruct transmission chains (Bryant *et al.*, 2013; Sabat *et al.*, 2013). The disadvantage using a single reference to compare different individuals is that features missing from the reference cannot be detected. Especially in the absence of a closely related reference genome, such an approach is not appropriate (Abdelbary *et al.*, 2018). To overcome this problem, an increasing number of studies incorporate the pan-genome of the species into the analysis of different isolates. Here, gene content and genomic rearrangements such as insertions, deletions, translocations, and inversions are used to explain the manifestation of phenotypic traits like antibiotic resistance (Medini *et al.*, 2005). One approach to compute a pan-genome is based on whole-genome alignments (WGAs) (see for example Angiuoli *et al.* (2011); Schatz *et al.* (2014); Hennig *et al.* (2015)). In comparison to a BLAST-based approach or variants of it which are employed to compute orthologous gene groups, the WGA-based approach has the advantage that for the identification of orthologous genes gene neighbourhood is taken into account. In addition, the pan-genome based on a WGA can be generalized to take also non-genic features into account. Further applications of WGAs encompass the identification of pathogenic genomic islands or reconstruction of phylogenomic trees (Chan and Ragan, 2013).

Runtimes of current state-of-the-art aligners that are capable of modeling genomic rearrangements, such as `progressiveMauve` (Darling *et al.*, 2010), are at least quadratic in the number of genomes. Thus, computation of WGAs of hundreds or thousands of even closely related bacterial genomes with state-of-the-art tools are prohibitive. Currently only `Parsnp` (Treangen *et al.*, 2014) is able to compute large-scale alignments with hundreds or even thousands of genomes within hours. However, it computes only a core-genome alignment. While `Parsnp` is a highly valuable tool for the identification of SNPs from the core pan-genome, it does not allow for the detection of genomic rearrangements and pathogenic islands for example. Recent advances in the field of whole-genome alignment methods were made with the introduction of `seq-seq-pan` (Jandrasits *et al.*, 2018), which offers the efficient extension of existing WGA by new genomes. Through a pairwise iterative alignment process, new genomic sequences are aligned against the consensus sequence of the alignment, which shows a substantial runtime decrease while achieving comparably high-quality alignments.

`seq-seq-pan` makes use of the profile of a pairwise alignment (and deduces a consensus sequence) for extension by new sequences. Using a profile as a representation for an alignment is a popular concept used to compute multiple sequence alignments (MSA) and was first introduced by Hogeweg and Hesper (1984) in their progressive alignment heuristic. The idea to progressively compute MSAs along a guide tree, where at each node either a pairwise alignment between sequences, sequence-profile or profiles are conducted, reduces the runtime significantly in contrast to the construction of an optimal alignment that maximizes the sum of pairs score. Especially the profile-profile alignment offers a fast and efficient way to combine two separate alignments. Recent advances in this field have been made by Liu and Warnow (2014) with the idea of a divide and conquer approach to compute smaller subsets of alignments which are merged through a profile-profile alignment. The parallel computation of the subset alignments further reduces the computational runtime, while still achieving highly accurate results.

However, currently such a profile-alignment based approach is missing in state-of-the-art alignment tools such as `progressiveMauve`. Since WGAs need to consider genomic rearrangements, such as translocations and inversions, between the individual genomes, a profile alignment of two or more WGAs is more difficult than a profile alignment of MSAs. Here, we introduce our concept for the first profile-profile alignment of WGAs. The profile-based merging of WGAs is conducted with the help of our SuperGenome data structure (Herbig *et al.*, 2012), which can be used to transfer different WGA-coordinate systems into a common one.

Based on this profile-based alignment approach, we implemented `GPA` (Genome Profile Alignment), a software that can align hundreds to thousands of genomes in a fast and efficient manner. The goals of our whole-genome alignment construction strategy were to adopt the advances from the field of profile-based MSA tools, and therefore significantly reduce computational time and still achieving highly accurate WGAs. Our intention was not to develop a new genome alignment algorithm. Therefore, `GPA` relies on other whole-genome aligners but using a divide and conquer strategy to merge subsets of alignments along a guide tree. In our current release, we have combined `progressiveMauve` with our profile-based approach.

The article is organized as follows: In the method section, we first present the SuperGenome data structure and the algorithmic principles that allows an efficient merging of several alignments. The critical aspect is the transfer of different coordinate systems into a common one, which is supported by the SuperGenome. We explain in detail the extension of a given WGA by other genomes or other WGAs. Based on this we then explain how to compute large-scale WGAs from scratch in a fast and parallelized manner. The section concludes with a description of statistics that we used to compare and evaluate our approach with the original `progressiveMauve`. The results section presents the evaluation of the WGAs computed for different simulated as well as real biological data sets by `progressiveMauve` and `GPA`. The evaluation is focused on the runtime needed and the achieved quality of the WGA by both approaches. Finally, we conclude this article with a critical discussion of our results and propose future extensions of our approach.

## 2 Methods

### SuperGenome data structure and construction

In comparison to multiple sequence alignment, where the order of nucleotides within each sequence is assumed to be preserved, aligning whole-genomes has to consider the occurrence of rearrangements, such as translocations as well as inversions. Regions shared by two or more genomes that do not contain any rearrangements of homologous sequences are called locally collinear blocks (LCBs), and a WGA is typically then represented by a set of such LCBs. Programs computing WGAs of this form are for example `Mauve` (Darling *et al.*, 2004), `progressiveMauve` (Darling *et al.*, 2010), `Mugsy` (Angiuoli and Salzberg, 2010) and `TBA` (Blanchette *et al.*, 2004). The SuperGenome is a data structure, which makes use of a whole-genome alignment (WGA) and features a bidirectional mapping between the alignment coordinates and the original coordinates of each individual genome in the WGA (Herbig *et al.*, 2012). The main advantage of the SuperGenome data structure is that it provides an unambiguous coordinate system and is independent of any pre-chosen reference genome.

We will first introduce some formal terminology before we then describe the algorithm to compute large-scale WGAs to decrease computational runtime using a profile-based approach together with the SuperGenome data structures derived from the profiles.

Given are *n* genomes *g*_*i*_, *i* = 1, …, *n* and a WGA **A** on these *n* genomes. We define a pair of integer arrays *G*_*i*_ and *SG*_*i*_, which provides the bidirectional mapping between the positions of *g*_*i*_ and **A**. Both arrays cover either all genomic or alignment positions and are zero-based numbered. In addition, the arrays have a leading entry, that is used to represent the absence of the sequence and are due to that one position longer than either the length of the genome or the alignment. This simplifies the mapping of the coordinate systems since the *j*-th index represents the *j*-th position in the genome or the alignment.

Let us assume that the *j*-th base of *g*_*i*_ is aligned at the *k*-th position in **A**. Thus, *G*_*i*_, representing the mapping of *g*_*i*_ to **A**, contains the value *k* at entry *j*. In the case that the base was aligned as its reverse complement (i.e., representing an inversion), the value of *k* is negative:

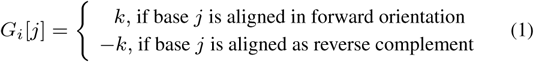

The respective SuperGenome aray *SG*_*i*_ represents the mapping of the aligned sequence of genome *g*_*i*_ in **A** and is analogous to *G*_*i*_, in addition, it also accounts for gaps. In case at position *k* in **A** no base of *g*_*i*_ was aligned the entry is set to zero.

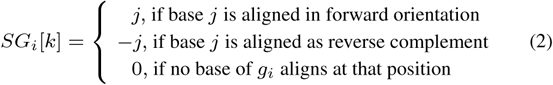

In addition to the coordinate mapping, the SuperGenome takes the local collinear block (LCB) structure into account, as for example, computed by `progressiveMauve` and stored in y`XMFA`-format, by also tracking the start and stop positions of all LCBs.

For a given WGA of *n* genomes, the generation of the data structure is straightforward. For each genome, the pair of arrays *G*_*i*_ and *SG*_*i*_ is initialized with the corresponding lengths of *g*_*i*_ and **A**, respectively, and filled with zeros. Through an iteration over every alignment position, the arrays are filled, resulting in a total of 2×*n* arrays with a space requirement of *n*× length of the alignment and the sum of all genome lengths. From the SuperGenome data structure, the WGA can be derived from the arrays *SG*_*i*_, where each aligned sequence of *g*_*i*_ can be reconstructed iteratively back from the genomic positions of the entries. Here, inversions (negative values) and gaps (zeroes) have to be taken into account.

### Extending an existing WGA

We now first describe how a new genome is added to an existing WGA on *n* genomes using the SuperGenome approach. Then we show how to extend this principle to merging two WGAs on *n* and *m* genomes, respectively, into a WGA on *n* + *m* genomes. The general idea is that for the extension step only a pairwise alignment is computed. This concept can then be generalized to several alignments and genomes that are combined into a WGA of all involved genomes.

A common approach to make use of an existing alignment **A** is a profile alignment, which preserves all prior aligned positions. The profile of **A** is represented by a consensus sequence, which is derived from the SuperGenome data structure through a majority call on all alignment positions of **A** (see Figure 1A for an example). For the integration of a new genome *g*_*n*_+1 into **A**, unique regions of *g*_*n*_+1 and homologous regions of *g*_*n*_+1 and the profile have to be computed in order to extend **A** by the *g*_*n*+1_. This is achieved by computing a pairwise alignment of the profile consensus sequence and the genomic sequence *g*_*n*_+1. This pairwise alignment serves as a guiding alignment to extend the given alignment **A** by the new genome *g*_*n*_+1. Again we use the SuperGenome data structure for this step (see Figure 1B-D for an illustration of this procedure). For this, the SuperGenome data structure of the pairwise alignment is computed, which includes the array *SGcons* with all positions of the profile consensus sequence and *SG*_*n*_+1, the SG array of genome *g*_*n*_+1. The extension of **A** comprises the transfer of the coordinates of *SG*_*i*_ and *G*_*i*_ for the *n* genomes into the common coordinate system of *SG*_cons_ and *SG*_*n*_+1. The new bidirectional coordinate mappings 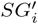 and 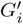 for genome *g*_*i*_ are derived by

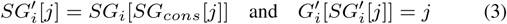

where *SG*_*cons*_[*j*] is the index of the consensus sequence, which is aligned at position *j* in the pairwise alignment, and *SG*_*i*_[*SG*_*cons*_[*j*]] the index of the base of *g*_*i*_ that was aligned in **A**.

**Fig. 1.**
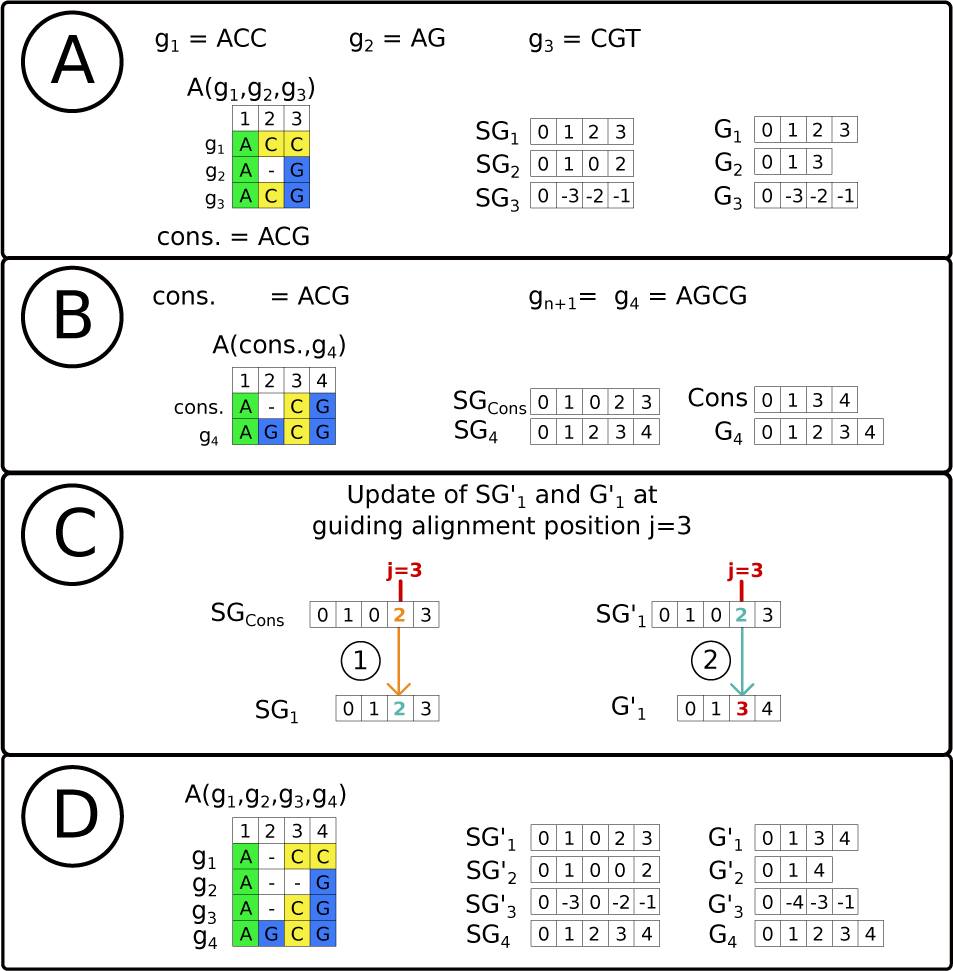
(A) Based on the three genomic sequences *g*_1_, *g*_2_ and *g*_3_ and their alignment **A**(*g*_1_, *g*_2_, *g*_3_), a SuperGenome data structure (union of the *SG* and *G* arrays) is computed. In addition, a consensus sequence from the alignment is deduced. (B) A new genomic sequence *g*_4_ is combined with **A**(*g*_1_, *g*_2_, *g*_3_). For this a pairwise alignment **A**(*cons*, *g*_4_) is computed and again a SuperGenome data structure is deduced. (C) Update of **A** by *g*_4_: for every position *j* in **A**(*cons, g*_4_), (1) array *SG*_*cons*_[*j*] (orange) contains the index of the aligned consensus sequence positions, which is used to determine the original genomic positions *SG*_1_ [*SG*_*cons*_[*j*]] (example shown in blue). (2) This allows a coordinate transfer 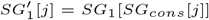 and 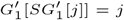 (red) into a common coordinate system of **A**(*cons*, *g*_4_). (D) From the updated SuperGenome data structure the new alignment **A**(*g*_1_, *g*_2_, *g*_3_, *g*_4_) is easily deduced.

This coordinate transfer restores all columns of alignment **A**. The arrays *SG*_*n*+1_ and *G*_*n*_+1 do not have to be updated since they are already consistent with the coordinate system of the guiding alignment. Based on the updated SuperGenome data structure on *n* + 1 genomes, the new alignment can now be easily derived and written into the alignment format as described above.

The procedure for adding one genome to a WGA is easily extended to merging the profiles of two WGAs on *n* > 1 and *m* > 1 genomes, respectively. Again, we compute a pairwise alignment, now on the two respective consensus sequences, derive the respective SuperGenome, which together with the SuperGenome of the input WGAs to update the bidirectional mappings (see equation (3)) the two WGAs. A consequence of this merging procedure is that prior aligned bases remain aligned in the merged WGA.

Finally, it is straightforward to generalize the pairwise approach to combining more than 2 WGAs or WGAs with individual genome sequences. Rather than aligning only two sequences, we compute the multiple genome alignment of all consensus sequences derived from the *k* > 2 individual WGAs as well as possible single genome sequences. The generalized workflow of merging several WGAs into a common WGA can be summarized in the following 6 steps (see also Figure 2):

**Fig. 2.**
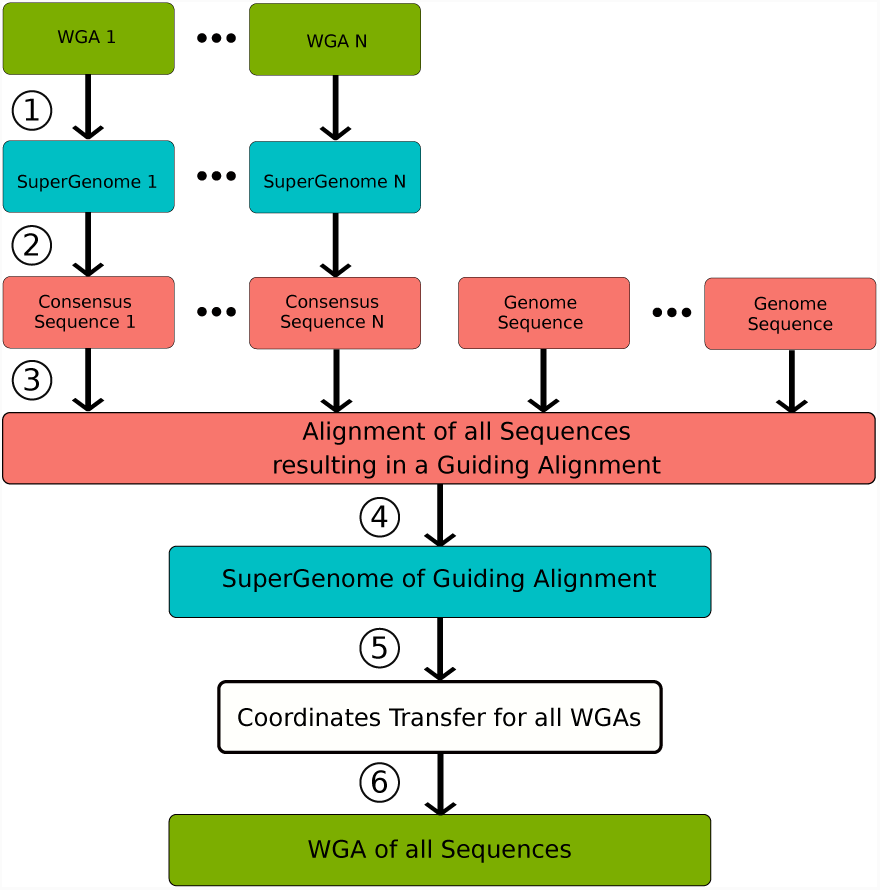
Workflow of merging several alignment and genomes using the SuperGenome data structure into a large WGA. The workflow consists of 6 steps: 1. Build SuperGenome for every alignment, 2. Compute consensus sequence from input alignments, 3. Align all consensus and genome sequences, 4. Build SuperGenome of guiding alignment, 5. Merge all alignments and genomes via coordinates transfer, 6. Output alignment of all sequences in XMFA-format derived from updated SuperGenome data structure.

1. Construct SuperGenome data structure for every input alignment.
2. For every SuperGenome compute a consensus sequence (output in FASTA format).
3. Align consensus sequences as well as possible individual genome sequences with whole-genome aligner (e.g., using `progressiveMauve`) to generate the guiding alignment.
4. Construct SuperGenome data structure for the guiding alignment.
5. For every genome *i*, update *SG*_*i*_ and *G*_*i*_ according to equation (3). Output new alignment derived from updated *SG′* and *G′* (output in `XMFA`-format).

The merged WGA also accounts for LCB structures from the input alignments. Previous start and stop positions, which determine the borders of a LCB, are transferred and added to the ones introduced by the guiding alignment. The resulting LCBs of the merged WGA are defined between every consecutive pair of LCB border position originating from the different WGAs that are merged to secure the LCB property that no rearrangements within a block occur.

### Genome Profile Alignment - GPA

We have implemented the described approach how to efficiently merge several whole-genome alignments or extend a given WGA by new genomes using our SuperGenome data structure in the tool which we call `GPA`. Our tool is written in Java and can be run on any machine with a Java VM installed. For the computation of the WGAs, we currently make use of `progressiveMauve`, which needs to be installed independently.

`GPA` can be applied in two ways: it can align genome sequences from scratch, or it can be used to extend an existing WGA by new genomes. In the first case, the input data are the genome sequences that need to be provided in FASTA-format. Since the general idea of our approach is to combine smaller sets of aligned genomes to a full WGA, these smaller sets first have to be defined. For this, we either make use of a guide tree, which determines the individual merging steps, or all genome sequences are randomly distributed into subsets, where the size of the subsets has to be predefined by the user. The guide tree needs to be provided by the user in Newick tree format and can, for example, be the one as computed by `progressiveMauve`. As most guide trees are binary and only two sequences are aligned at each node, the provided input tree is further modified, to control the number of sequences/WGAs that are aligned in each step. With a user-defined maximum number of sequences aligned in each step, the nodes of the guide tree (representing the set of sequences which will be aligned) are propagated from the leaves (representing the genomes) towards the root (representing the final WGA) until another propagation to the next node of the tree would exceed the maximum. This modified guide tree is used to compute an internal guide tree structure. In the next step, after the internal guide tree structure has been built, `GPA` automatically creates a folder structure, which serves to save the WGAs from the intermediate steps in `XMFA`-format. The computation of the WGAs follows the typical process of progressive alignments, starting at the leaves of the guide tree and the root represents the full WGA. If no guide tree has been provided, `GPA` merges all WGAs of the individual subsets into a common WGA in one step. To further decrease the runtime, `GPA` provides the possibility to compute the independent subalignments in parallel.

The second application case is to extend an existing WGA. For this, `GPA` can be provided with an arbitrary number of WGAs and single genomes. Here, the respective profiles or genomic sequences are aligned in one step. The final WGA then contains all new genomes as well as those which where contained in the input WGAs. Note that in fact the second case is used throughout the computation of a WGA from scratch when using our approach in `GPA`.

### Experimental setup

#### Evaluation Criteria

Besides runtime assessment, we used three different statistics to compare the whole-genome alignment computed using our guide-tree based SuperGenome approach with the whole-genome alignment computed by applying `progressiveMauve` to all genomes at once: *pairwise consistency score*, *total column score*, and *F-score*.

The pairwise consistency (PC) score reflects how much the WGA agrees with all possible pairwise alignments, a concept which was first described by Gotoh (1990) and adapted in `T-COFFEE` (Notredame *et al.*, 2000). For this, for each pair of genomes in the WGA we compute the percentage of bases that are consistently aligned in the WGA and in the respective pairwise genome alignment. For a given WGA of *n* genomes, we then report the average pairwise consistency score from all 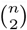 scores.

The total column (TC) score, first introduced in `BaliBase` (Thompson *et al.*, 2005), equals the percentage of identically aligned columns in a given alignment when compared to a so-called reference alignment.

The third statistic, the *F*-score, is the harmonic mean of *precision* and *recall* for identical pairwise aligned bases. The *F*-score was used in the Alignathon competition (https://compbio.soe.ucsc.edu/alignathon/) for the comparison of two WGAs (Earl *et al.*, 2014), where one of the two WGA serves as a reference.

All three statistics have been widely used to evaluate multiple sequence alignment methods. The total column score is very conservative and prone to small changes in the alignment, since one misaligned sequence can drastically decrease the score. Therefore, comparing the best multiple sequence alignment methods may only result in low TC scores. On the other hand, only slight differences in the PC score between two alignments, as well as a high TC and *F*-score, are reliable indicators for the similarity of the two. For the calculation of all scores when comparing two WGAs, one needs to take care of possible inversions and translocations. Again, the SuperGenome data structure serves extremely useful for this, and therefore we also used it for the calculation of these scores.

#### Simulated data sets

To evaluate the performance of `GPA` in the context of a ‘ground truth’, where the optimal WGA is known, we used the same genome simulation approach as the Alignathon. The simulated genomes were generated with the software `EVOLVER` (Edgar *et al.*, 2009) together with the `evolverSimControl suite` (Edgar *et al.*, 2011), that is able to evolve genomes based on a given phylogenetic tree and report the respective WGA. The simulation parameters required for the `evolverSimControl suite` were used from the `EVOLVER` example and adapted to prohibit duplication events, since Mauve cannot address duplication events. The *F*-scores between the simulated and `GPA` generated WGAs were calculated with `mafTools` (Earl *et al.*, 2014) (https://github.com/dentearl/mafTools/).

#### Biological data sets

To explore the performance of `GPA`, we applied our approach to a large number of complete single chromosome bacterial genomes from the same species which we derived from NCBI (ftp://ftp.ncbi.nlm.nih.gov/genomes). The various data sets reflect different genome lengths and sizes as well as diversities of genomes within a species to explore the performance of `GPA`. Currently, `GPA` can only handle single sequences per genome. However, it is possible to concatenate the different chromosomes (or plasmids) of one genome to overcome this limitation. Still, we decided to remove all possible plasmids prior to the WGA computation and only computed single chromosome alignments. All data sets used in this work are listed in Table 1 together with the total number of genomes as well as the average genome length and average GC content (Source https://www.ncbi.nlm.nih.gov/).

**Table 1.** The data sets, which were used for the WGA computations as obtained from the NCBI FTP server. All statistics including the median genome length and median GC content have been derived from NCBI.

| Organism | # Strains | Median Genome Length (Mb) | GC content |
| --- | --- | --- | --- |
| *Bacillus cereus* | 13 | 5.760 | 35.1% |
| *Listeria monocytogenes* | 30 | 2.975 | 37.9% |
| *Chlamydia trachomatis* | 72 | 1.046 | 41.3% |
| *Mycobacterium tuberculosis* | 128 | 4.385 | 65.6% |
| *Klebsiella pneumoniae* | 166 | 5.590 | 57.2% |
| *Staphylococcus aureus* | 176 | 2.847 | 32.8% |
| *Bordetella pertussis* | 326 | 4.100 | 67.7% |

#### Computational Platform

We ran all WGA computations on a Linux server with four *Intel(R) 197 Xeon(R) CPU E5-4610 v2 @ 2.30GHz* and 500 GB of memory. We measured the runtime with the `GNU time` command. During all WGA computations, `GPA` used the maximal number of threads needed to ensure that all independent subset alignments could be computed in parallel.

## 3 Results

With `GPA` we have extended `progressiveMauve` by the possibility to provide an existing sequence alignment in `XMFA`-format and align it to other sequences or alignments. Traditionally, profile alignment is conducted on a pairwise level, however with `GPA` a multiple profile alignment can be computed in one step. With this feature, a progressive alignment strategy that is typically performed along a binary guide tree can now be generalized to non-binary trees with fewer internal nodes. `GPA` provides a parameter *k*, that controls the maximal degree of multifurcation of every internal node and therefore how many genomes and/or profiles are merged at each step. Our overall intention for this strategy was to multiply align up to hundreds or more bacterial genomes with a significantly reduced runtime and at the same time achieve highly qualitative WGAs. We first evaluated the WGAs computed by `GPA` based on simulations, followed by comparison of the WGAs computed for real biological data sets.

### Evaluation based on simulated data

We simulated five different WGAs of different sizes, where we used the genome of *B. pertussis* (RefSeq ID: *NC_002929.2*) as the start of the simulation as well as the guide trees computed from real biological genomes (see below) using `progressiveMauve`. The simulations were all independent of each other. Therefore a comparison of the PC, TC and F-Scores across the WGAs of different size is misleading because only the phylogenetic origin of the simulated data set is same. Here, the focus is set on comparing the performance of the iterative profile alignment approach of `GPA` with `progressiveMauve`. For `GPA`, WGAs with different merge sizes *k* were computed, the one with the highest *F*-score is reported.

As can be seen from the results (see Table 2) the computation of the WGA by `GPA` is not only significantly faster than computation by `progressiveMauve`, but also the runtime difference increases with increasing number of genomes. At the same time the WGA computed by `GPA` is highly comparable to the one computed by `progressiveMauve`. For example, both WGA computation approaches yield highly similar PC scores, which differ in less than 2% (data for different *k* is not shown). Also, the combination of high TC (> 60%) and *F*-scores (> 0.98) between the respective WGAs of both tools shows their high conformity. In addition, the *F*-scores, derived from the comparison of the WGA as computed by `progressiveMauve` and `GPA` respectively with the simulated, ‘ground truth’ WGA are almost identical. The largest difference between both approaches is the memory needed to construct the alignment. Here, `GPA` needs around 3 to 5 times more memory than `progressiveMauve` to compute the WGAs. An exception for this is WGA computation of 80 simulated strains, where the increase of memory consumption as well as runtime of `progressiveMauve` is extraordinary. The simulation was repeated several times, and the large runtime and memory consumption was confirmed for each run.

**Table 2.**
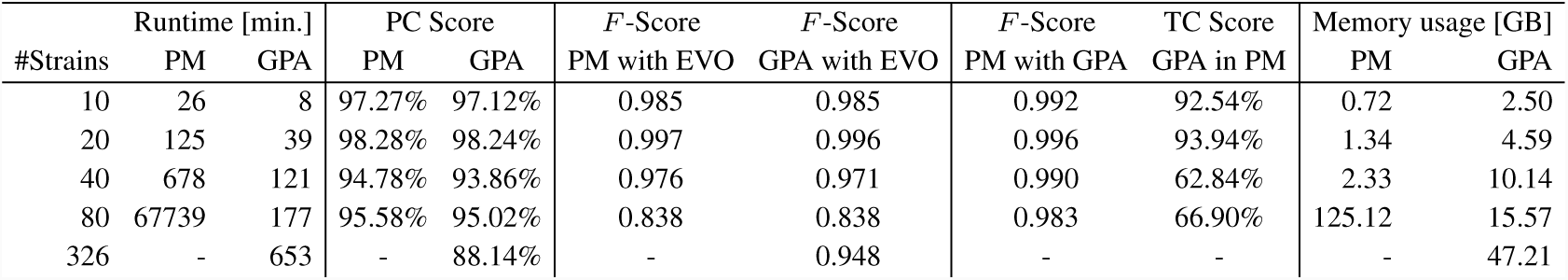
Evaluation of WGAs generated by `progressiveMauve` (PM) and `GPA` based on simulated WGAs by `EVOLVER` (EVO). All WGAs were evaluated with respect to their runtime, average pairwise consistency (PC), *F*-scores achieved against EVO and maximal amount of RAM used in the process of WGA computation. In addition the total column (TC) score and *F*-score between PM and `GPA` is reported. Again, for the calculation of the TC and *F*-score, PM is used as the reference. `GPA` was run with several different parameters *k* for the merge size, reported for each data set is the one with the highest *F*-score achieved with EVO.

### Evaluation based on biological data: Runtime

Since, in the current implementation of `GPA` we use `progressiveMauve` as underlying multiple genome alignment method, our evaluation focuses on the direct comparison between the WGA produced by applying `progressiveMauve` to all genomes at once and the WGA computed using our iterative merging approach implemented in `GPA`.

To compare the runtimes for the WGA construction of both `progressiveMauve` and `GPA`, all WGAs are computed from scratch. Here, for data sets with less than 100 genomes we chose to split these into randomly distributed groups that were of equal size if possible. For all other data sets we used the guide tree produced by `progressiveMauve`, to determine the individual merging steps. In addition, for data sets with more than 100 genomes available, we computed several WGAs of different size. Starting at a set of 10 genomes, we subsequently added new genomes to the set for the next larger alignment. This ensures that we can evaluate the progression of the performance of the tools with the increasing number of genomes.

The results (see Table 3) show a general significant runtime decrease for the WGA construction of `GPA` compared to `progressiveMauve`, independent of the data set. As it can be seen from the results and Figure 3, the runtime of `progressiveMauve` increases at least quadratically with the number of aligned genomes, while for `GPA` the increase shows a linear dependency. The scalability and runtime reduction of `GPA` is most impressive for the WGAs with over 80 genomes. Here, none of the WGAs computed by `progressiveMauve` have finished after 350 hours or even 1650 hours (more than two months) of computing time, where the choice was made to not further wait for the result. Note in all these cases, the guide tree was still constructed. Furthermore, `progressiveMauve` did not finish the WGA with 80 strains of *K. pneumonia* within 1400 hours. Therefore the run was aborted, the respective results are also not stated and compared to `GPA`. In contrast to this, `GPA` could compute all WGA computations within a couple of hours or maximally few days. For example for the WGA of 176 *S. aureus* strains `GPA` needed slightly more than 3 hours (198 minutes), while `progressiveMauve` ran more than 1650 hours without reporting a result, thus in this case `GPA` was at least 500 times faster.

**Table 3.** Comparison between the WGA construction using the original `progressiveMauve` (PM) method and our SuperGenome-based iterative profile alignment approach of `GPA`. The results are divided into two distinct groups, whether for `GPA` a guide tree (guide tree) was used or not (random). The runtime for WGA computation and the number of LCBs for the respective WGAs is reported. In addition, the WGAs were evaluated with respect to their average pairwise consistency (PC), the total column (TC) score (% of identical aligned columns in PM) and *F*-score. Both, for the calculation of the TC and *F*-score, PM is used as the reference. `GPA` was run with several different parameters *k* for the merge size, reported for each data set is the one with the highest PC score.

| Data set | Organism | # Strains | Runtime [min] |  | # LCBs |  | PC Score |  | TC Score | *F*-Score |
| --- | --- | --- | --- | --- | --- | --- | --- | --- | --- | --- |
|  |  |  | PM | GPA | PM | GPA | PM | GPA |  |  |
| random | *B. cereus*<br>(5.760Mb) | 13 | 225 | 61 | 380 | 214 | 60.00% | 60.63% | 62.40% | 0.977 |
|  | *L. monocytogenes*<br>(2.975Mb) | 30 | 628 | 28 | 293 | 254 | 73.41% | 73.92% | 61.50% | 0.989 |
|  | *C. trachomatis*<br>(1.046Mb) | 72 | 351 | 14 | 4 | 80 | 98.35% | 98.36% | 86.68% | 0.995 |
| guide tree | *M. tuberculosis*<br>(4.385Mb) | 10 | 27 | 9 | 3 | 6 | 97.90% | 97.97% | 93.99% | 0.998 |
|  |  | 20 | 98 | 22 | 16 | 4 | 96.83% | 97.29% | 88.31% | 0.994 |
|  |  | 40 | 475 | 41 | 230 | 72 | 96.00% | 96.61% | 85.05% | 0.993 |
|  |  | 80 | 1665 | 105 | 852 | 148 | 92.36% | 93.03% | 73.03% | 0.990 |
|  |  | 128 | >350h* | 527 | - | 437 | - | 90.67% | - | - |
|  | *K. pneumoniae*<br>(5.590Mb) | 10 | 55 | 25 | 177 | 160 | 38.77% | 38.94% | 73.13% | 0.991 |
|  |  | 20 | 691 | 90 | 784 | 769 | 41.31% | 41.42% | 67.18% | 0.989 |
|  |  | 40 | 9401 | 99 | 3369 | 1463 | 39.32% | 43.15% | 55.94% | 0.981 |
|  |  | 80 | >1400h* | 329 | - | 4632 | - | 42.38% | - | - |
|  |  | 166 | >6250h* | 771 | - | 8558 | - | 41.19% | - | - |
|  | *S. aureus*<br>(2.847Mb) | 10 | 16 | 7 | 221 | 53 | 82.45% | 81.81% | 73.07% | 0.983 |
|  |  | 20 | 64 | 33 | 415 | 362 | 81.58% | 81.11% | 67.83% | 0.986 |
|  |  | 40 | 552 | 28 | 1222 | 555 | 76.33% | 75.70% | 56.63% | 0.976 |
|  |  | 80 | 5213 | 121 | 2492 | 2498 | 73.02% | 72.50% | 47.37% | 0.976 |
|  |  | 176 | >1650h* | 198 | - | 3213 | - | 71.91% | - | - |
|  | *B. pertussis*<br>(4.100Mb) | 10 | 24 | 9 | 64 | 56 | 58.77% | 58.99% | 95.87% | 0.995 |
|  |  | 20 | 99 | 16 | 88 | 115 | 58.56% | 58.45% | 90.06% | 0.991 |
|  |  | 40 | 503 | 96 | 165 | 172 | 57.65% | 57.78% | 85.48% | 0.990 |
|  |  | 80 | 1683 | 78 | 314 | 373 | 58.69% | 58.39% | 75.57% | 0.981 |
|  |  | 326 | >6250h* | 504 | - | 3708 | - | 47.37% | - | - |

\* time past till computation has been manually been aborted by us

**Fig. 3.**
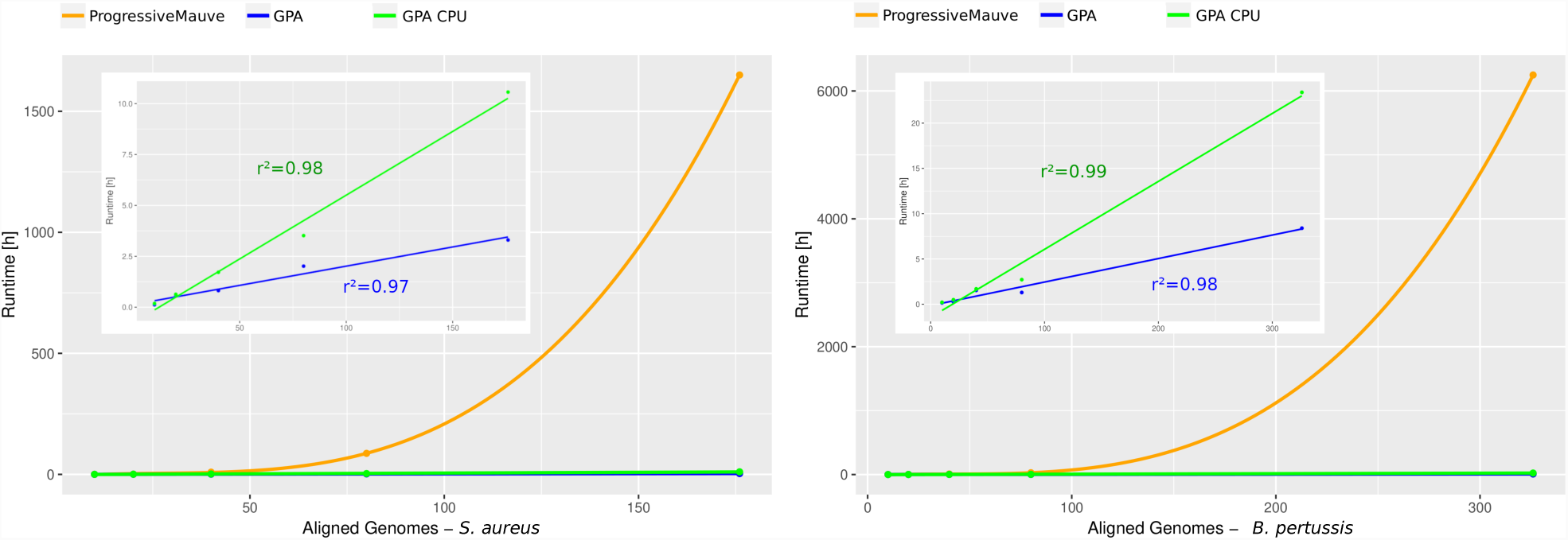
Comparison of the measured computational runtime needed for the construction of the WGA depending on the number of genomes for the data sets of *S. aureus* (left) and *B. pertussis* (right). In addition to the direct comparison between `progressiveMauve` (orange) and `GPA`, the upper left section only shows the runtime of `GPA` (blue) and `GPA` CPU time (green), together with the *r*^2^ values for the linear regression.

Independent from the used WGA construction method, the runtime comparison between the data sets of the species show differences as well. Here, an essential factor is the average genomic length. The computation time needed for WGAs with the same number of aligned genomes was less for the shorter genomes of *S. aureus* than for *M. tuberculosis* and *B. pertussis*, which show comparable average genome lengths as well as runtimes. The WGAs for *K. pneumonia* that has on average the longest genomes also needed the longest absolute runtime. When comparing the results of *M. tuberculosis* and *B. pertussis*, which have a similar length, the diversity of the genomes within a data set (reflected by a smaller overall pairwise consistency score) does not affect the runtime as much as the genome length.

The number of LCBs, which represent genomic architectural differences between the genomes, differ a lot between the organisms analysed here. For example, WGAs of *M. tuberculosis* and *B. pertussis* with similar genome lengths have vastly different number of LCBs. On the other hand, as can be seen from Table 3 with increasing number of aligned genomes from the same organism also the number of LCBs always increases, independent whether the WGA was computed using `progressiveMauve` or `GPA`. When comparing `progressiveMauve` and `GPA` for a given set of genomes, the number of LCBs is largely of similar order of magnitude, though in most cases (but not all) the WGA computed with `GPA` had less LCBs than those for the WGA computed with the original `progressiveMauve`.

### Evaluation based on biological data: Qualitative comparison of WGAs

In order to compare the WGA computed using `GPA` with the one computed by the original approach of `progressiveMauve`, we applied three evaluation criteria (see Methods section). For each data set, we calculated the average pairwise consistency (PC) score, total column (TC) score and *F*-score (see Table 3).

For each data set, independent of the number of aligned strains per species, the average PC score is very similar between the `progressiveMauve`- and `GPA`-derived WGAs (less than 1% difference). The largest difference of 3.8% is observed for the alignment of 40 strains of *K. pneumoniae*, where `GPA` yields a higher PC score than `progressiveMauve`. Overall, the pairwise consistency only slightly decreases within a species when adding more genomes to the alignment. The largest difference was observed for the WGAs computed for *B. pertussis*: here the average PC score dropped from 58.99% for 10 genomes down to 47.37% when 326 genomes were aligned. An exception has been observed for the *K. pneumoniae* WGAs. Here the average PC score of the WGA built from 10 and 20 genomes was smaller than for the WGAs with 40 and 80 genomes. Generally, the PC score differs most strongly when comparing different bacteria. While the WGAs of *M. tuberculosis* and *C. trachomatis* have average PC scores greater than 90%, the WGAs of *K. pneumoniae* achieve a maximum of 43%.

The TC score, which represents the fraction of identically aligned columns, compares the WGAs computed by `GPA` with the one derived with the original `progressiveMauve`. Overall the TC score is above 60% in most cases, indicating that `GPA` aligns a majority of all columns identically to the original `progressiveMauve` even for larger WGAs with up to 80 genomes. Also, none of the WGAs of our test data sets shows a TC score below 45%. Similar to the PC score the TC score differs between different organisms though here the biggest differences are seen when increasing the number of genomes within a species. Interestingly, the PC and TC scores do not necessarily correlate a lot, i.e., WGAs with similar PC scores do not necessarily have similar TC scores. For example, WGAs with low PC scores (as seen for example in the case of *B. pertussis*) may have higher TC scores than WGAs with high PC scores (e.g. *S. aureus*).

Another indicator of the high similarity of the WGAs computed with `GPA` and those computed with `progressiveMauve` is the very high *F*-score. Independent of the organism as well as the number of genomes the smallest value is 97.6% and the largest value is 99.8%. Here, in general the *precision* score is higher than the *recall* for the resulting *F*-score (data not shown). We observed that WGAs with a high TC score also have a high *F*-score. On the other hand, increasing the number of genomes for a WGA generally leads to a significant decrease of the TC score, while this behavior is not observed for the *F*-score. An example is *S. aureus*, where the lowest TC score in all comparisons was achieved, while the *F*-score is still above 0.97.

### Impact of compressing the guide tree

Next, we analysed the impact of the multifurcation parameter *k*, which is used to compress the input guide tree. This parameter *k* reflects the maximal degree of each internal node of the guide tree, which represents the maximal number of profiles or sequences that are merged in a single step during the WGA computation. Clearly, as can be seen from Table 3, the runtime of `progressiveMauve` increases significantly with increasing genomes. Thus, a trade-off between the number of genomes aligned at a time and runtime needs to be considered when choosing *k*.

For this purpose, for a given initial guide tree on a given set of genomes, we computed WGAs with `GPA` using different *k*. For each *k* we reported the runtime, PC, TC as well as the *F*-score. As expected larger *k* result in larger runtimes (see Table 4 for the case of *M. tuberculosis*). On the other hand, the PC and TC score, as well as the F-score, are not greatly affected by the choice of *k*, while the number of LCBs varies, though a clear pattern cannot be deduced. Similar observations have also been made for the other data sets of *S. aureus*, *K. pneumoniae* and *B. pertussis* (data not shown). Furthermore, the analysis of the WGAs, where no guide tree was provided for `GPA`, and therefore equally sized groups were merged, shows that at least for smaller sized WGAs highly similar results to `progressiveMauve` can be achieved as well (see upper parts of Table 3 and 3).

**Table 4.** Evaluation of `GPA`-derived WGAs with respect to the maximal number of sequences merged at a time for the *M. tuberculosis* data set.

| #Strains | max. degree<br>*k* | Runtime<br>[min.] | PC<br>score | TC<br>score | #LCB | *F*-Score |
| --- | --- | --- | --- | --- | --- | --- |
| 10 | 3 | 6 | 97.97% | 93.99% | 6 | 0.997 |
| 10 | 5 | 8 | 97.78% | 93.59% | 8 | 0.996 |
| 10 | 7 | 10 | 97.77% | 93.75% | 4 | 0.997 |
| 20 | 3 | 9 | 97.25% | 87.58% | 16 | 0.993 |
| 20 | 5 | 13 | 97.13% | 87.57% | 11 | 0.992 |
| 20 | 7 | 15 | 97.29% | 88.31% | 4 | 0.994 |
| 20 | 9 | 22 | 97.26% | 87.93% | 32 | 0.993 |
| 40 | 7 | 32 | 96.39% | 84.78% | 69 | 0.992 |
| 40 | 9 | 36 | 96.61% | 85.05% | 72 | 0.993 |
| 40 | 12 | 66 | 96.33% | 84.52% | 138 | 0.992 |
| 80 | 12 | 78 | 92.70% | 72.10% | 249 | 0.989 |
| 80 | 17 | 101 | 93.03% | 73.03% | 148 | 0.990 |
| 80 | 22 | 241 | 92.59% | 73.01% | 404 | 0.990 |
| 128 | 17 | 177 | 90.41% | - | 475 | - |
| 128 | 22 | 343 | 90.67% | - | 437 | - |

## 4 Discussion

In this paper we introduced `GPA`, a tool which extends the whole-genome alignment method `progressiveMauve` (Darling *et al.*, 2010) by the possibility to align individual genomes or whole-genome alignments to a given profile genome alignment. The challenge in the case of profile-based WGA is that genomic rearrangements like translocations and inversions have to be considered and therefore the profile of genome alignments cannot be as easily derived as in typical multiple sequence alignments. Each individual genome as well as whole-genome alignment represents its own coordinate system, and one challenge when merging a given WGA with individual genomes or another WGA is the transfer of the individual coordinate systems into a common one. With our SuperGenome data structure we have introduced a novel concept that allows the extension of a given WGA by further genomes or other WGAs, through a coordinate transfer along a guiding alignment of their profiles. Currently, we use `progressiveMauve` together with `GPA` and guarantee to adhere also to the locally collinear blocks computed by `progressiveMauve`.

Clearly, `progressiveMauve` has been shown to be among the best methods for the alignment of bacterial genomes because of its ability to consider rearrangements between the genomes. However, the runtime of `progressiveMauve` which was shown to be at least quadratic (Darling *et al.*, 2010), as well as lack of parallelization, prevent the computation of WGAs of hundreds or even thousands of genomes. Thus the second goal of this paper was to significantly reduce the runtime of `progressiveMauve`.

Though by definition the SuperGenome can be derived from whole-genome alignments of arbitrary species, we have restricted our analyses to intraspecies microbial genomes, i.e., aligning different strains from the same species.

One central feature of `progressiveMauve` is a binary guide tree and a WGA that is progressively computed along this tree. The key algorithmic ingredient in `GPA` is the use of a compressed rather than strictly binary guide tree and the adaptation of our introduced profile WGA to the alignment of several profiles in one step. The compressed guide tree leads to a significant reduction of internal nodes and therefore subalignments that are merged at a time. Using our SuperGenome based profile alignment approach together with a compressed guide tree, we could show that `GPA` is orders of magnitude faster than the original `progressiveMauve`. For example, for our largest dataset of *B. pertussis* with 326 genomes, we could show that `GPA` needed less than 9 hours, while a sole `progressiveMauve`-based alignment was not finished after 6250 hours (after which we aborted the computation). A linear regression analysis showed a clear linear relationship between runtime and genomes (see Figure 3). Based on the coefficients of the regression the computation of a WGA with 1000 *S. aureus* genomes for example would need about 19 hours, a WGA of 1000 *B. pertussis* genomes about 26 hours.

The results of `GPA` depend on the guide tree and the number of profiles that are merged at each node, our parameter *k*. As the guide tree determines the order in which the genomes are subsequently added, it is only plausible that the quality of the final WGA decreases if the tree does not reflect their phylogenetic relationship.

The comparison of `progressiveMauve` and `GPA` shows that as a trade-off for the reduced runtime of `GPA` the quality of the WGAs is in most of the cases slightly worse. However, the small differences of the pairwise consistency scores by only a few percents together with the high TC and *F*-scores show that the WGAs computed with `GPA` are highly similar to those computed using the original `progressiveMauve` approach. In addition, the comparison and evaluation of the simulated data shows, that when comparing the WGA computed by `progressiveMauve` and the one computed by `GPA` with the ‘ground truth’ both the PC score as well as *F*-score are almost identical independent of the number of genomes in the WGA. This shows that we could introduce a whole-genome alignment approach that enables a faster computation of WGAs, while achieving a comparable quality, which was our intention.

We also showed that there is a trade-off between runtime and quality when deciding how many profiles are merged at the same time. The parameter *k* controls the maximal number of profiles aligned at a time with `progressiveMauve`. On the other hand, larger *k* can lead to higher PC scores at the cost of increasing runtime. Since the results of `progressiveMauve` show that the computation time needed for WGAs with less than 20 genomes is in most cases below 2 hours, a *k* between 10 and 20 is currently our default and recommended value for larger WGAs with more than 50 genomes. In addition, the generation of WGAs with thousands of genomes by `GPA` benefits from the usage of large cluster systems, because the independent alignments can be computed in parallel.

Currently, `GPA` makes use of `progressiveMauve` as the underlying whole-genome alignment method, however, the modular implementation allows in principle the support of other whole-genome alignment methods. Since the SuperGenome data structure requires a one-to-one mapping between the nucleotides of the genomes, only methods are feasible that generate alignments with this feature. The disadvantage of these type of methods, which include `progressiveMauve`, is that duplicated regions cannot be aligned.

In the current version memory consumption is an issue of `GPA`, since the complete SuperGenome data structure is kept in memory to speed up computation. For example, for the alignment of 326 *B. pertussis* strains, `GPA` needed between 70GB and 110 GB RAM, depending on the parameter *k*. For the next release we plan to introduce an index file for the SuperGenome data structure that is accessed on demand to enable even larger WGAs with significantly less memory requirements.

The current version of `GPA` and the SuperGenome data structure uses the consensus sequence derived from the profiles of WGAs, which we used as a simple solution to efficiently align several profiles. Apparently, this is a feasible, but not an optimal solution, and therefore other methods that use a more sophisticated representation of the profiles for the merging step will be considered as possibly improved solutions in a future release. Furthermore, considering recent efforts with respect to the computation of optimal LCBs and their ordering (Gärtner *et al.*, 2018), which influence the structure of the coordinate system, could further improve our approach.

A possible application of `GPA` can be seen from comparative genome analyses that rely on a WGA. An example is AureoWiki (Fuchs *et al.*, 2017) (http://aureowiki.med.uni-greifswald.de/), a database of 32 different *S. aureus* strains that has been built from the results of a WGA-based pan-genome computation. To incorporate new *S. aureus* strains into the database, would require a WGA with the new genomic sequences. Computing this WGA from scratch could introduce changes that are not consistent with prior results. Here, the profile-based extension of the WGA by `GPA` preserves the former WGA and allows the addition of new strains without the necessity to completely rebuild the database.

As a conclusion, `GPA` introduces a time efficient computation of large-scale WGAs through the usage of WGA-profile alignments and adds further utility by the possibility to extend existing WGAs. With `GPA` hundreds to thousands or more bacterial genomes can now be fully aligned in an acceptable time.

